# Rapid Online Changes of Mind during Value-based Action Decisions

**DOI:** 10.1101/656330

**Authors:** Angela Martí-Marca, Gustavo Deco, Ignasi Cos

**Affiliations:** Center for Brain & Cognition (CBC), Pompeu Fabra University, Edifici Mercè Rodoreda, Carrer Trias i Fargas 25-27, 08005 Barcelona, Catalonia, Spain; Institució Catalana de Recerca i Estudis Avançats (ICREA), Passeig Lluis Companys 23, 08010 Barcelona, Catalonia, Spain

## Abstract

While the principles of decision-making are often expressed in terms of benefit-cost trade-off, reasonable doubt remains as to how the cognitive value and motor cost associated to several options may be computed across several brain areas, ultimately leading to a decision. Furthermore, does the assessment of non-chosen options continue after the decision has been made and the selected movement is already ongoing? Does the planning of several motor options depend on a unique parallel process or are there some elements within the sensorimotor loop requiring a sequential processing? Our hypothesis is that a change of available prospects at any point in time should dynamically adjust the desirability for each option, implying a fast reassessment of values and costs across prefrontal and motor cortical areas, even after an initial decision has been made and its associated response movement is ongoing. To test this, we performed a decision-making task in which human participants were instructed to select a reaching path trajectory from an origin to a wide rectangular target to gain the most reward. Reward delivery was contingent upon the distribution of value and upon precise target arrival. The original value distribution was altered during the ongoing movement in one third of the trials. Our results show that participants changed their movement trajectories towards the target position offering a better prospect, as presented by the second distribution. The changes of mind occurred more frequently for slow movements, had a duration in average inferior to the reaction time, and altered the initial timing and movement velocity. Although reward is the main factor guiding the selection of a specific movement in our experiment, our results indicate that the motor system was biased towards early changes of mind, given that the amplitude of the first acceleration and velocity peaks was significantly smaller in trials in which the participant switched target side and those in which they stuck to their original choice. Finally, the short latency of the recorded changes of mind strongly supports the hypothesis that, for the experimental conditions hereby considered, value considerations occur in parallel to motor planning.

## INTRODUCTION

Neural recordings have revealed that areas of the pre-motor cortex may simultaneously represent several option movements during the delay period of decision-making tasks (McPeek and Keller 2002; Cisek 2007; Cisek and Kalaska 2010) and that decisions ensue neural competition (Cisek and Kalaska 2005; Hikosaka et al. 1999). However, questions as to whether multiple actions continue being represented across the fronto-parietal loop after movement initiation, and how non-selected actions interact with the ongoing motor plan remain unclear. Previous evidence showed that situations of high sensory ambiguity facilitate changes of mind in visual decision-making (Resulaj et al., 2009), and that sudden perturbations to the state of the motor apparatus after the onset of movement may also result in a change of mind (Nashed et al. 2014). However, it remains to be established whether and how changes of mind could be voluntarily enacted during decision-making tasks when the distribution of reward associated to each choice changes after a decision has been made and the resulting movement is ongoing. Would these changes abide by the same cost/benefit trade-off observed in decisions between movements? (Stevens et al. 2005; Shadmehr et al. 2010; Rigoux and Guigon 2012). Since reward prospect is a major incentive modulating intentionality and vigour of movement (Choi et al. 2014; Mazzoni et al. 2007; Niv et al. 2007), it seems reasonable to hypothesize that the neural encoding of the desirability for each action should be sensitive to changes of value prospect within the environment, and adapt their assessment of associated cost and benefits accordingly across the fronto-parietal loop (Gold and Shadlen 2007; Kennerley et al. 2009; Wallis and Kennerley 2011; Padoa-Schioppa 2013; Thura and Cisek 2014), ultimately altering current movement trajectories to increase reward. To test this in a principled fashion, here we performed a decision-making task in which human participants selected and performed a reaching-path trajectory from an origin to a wide rectangular target. Reward was made contingent upon arrival precision by using bi-modal distributions around the right and left extremes of the rectangle in one out of three possible distributions. To test the influence of reward onto ongoing trajectories we changed the original distribution at different times after movement onset on approximately one third of trials. Our main results show that participants altered their initially selected path trajectory only when a better prospect was offered, according to the second distribution, along a path different to their original choice. Furthermore, changes of mind were more frequent for slow movements, involved an average duration inferior to the reaction time, and altered movement velocity. Although further research will be required to assess this, all of this is strongly supporting the notion that parallel planning and decision-making between motor actions, also when the valuation system is the main drive underlying decision-making.

## MATERIALS AND METHODS

### Participants

Twenty-three right-handed participants (8 males and 15 females, Mage = 24.0 yrs, SD = 5.0), participated in this study. Three participants (2F, 1M) did not comply with task instructions and their data was excluded from further analysis. The data from five other participants (3F, 2M) was corrupted, and was also excluded. A final sample of fifteen right-handed participants (6M and 11F; Mage = 24.4yrs, SD = 5.8) was analyzed. All had normal or corrected-to-normal vision and hearing, and did not suffer from any known neurological disorders. Informed consent was obtained following the guidelines established by the local ethics committee and all participants received monetary compensation (20€/hr) for their participation, regardless of completion.

### Materials and Experimental Setup

#### Equipment

Participants were seated in front of an Acer G245HQ computer screen (1920×1080) attached to an Intel i5 (3.20GHz, 64-bit OS, 4 GB RAM) computer. A spherical marker was placed on the tip of the right-hand index finger of subjects 1-12. Their movements were tracked with an Optitrak 3D motion tracking system (Optitrack, Inc). The motor trajectories of subjects 13-23 were recorded by sliding the computer mouse. All participants executed their motion trajectories while maintaining contact, at all times, with a flat, horizontal surface, in this case a table at a comfortable height in front of them. A chin-rest was used to stabilize posture at a fixed distance from the screen. The task flow was controlled with custom-built scripts, programmed using OpenFrameworks v.0.9.8. Data analyses were performed with custom-built MATLAB scripts (The Mathworks, Natick, MA), licensed to the Pompeu Fabra University.

### Experimental Task

Each experimental session consisted of a total of 630 trials. During each trial, the participant executed a free reaching movement from a circular origin cue (diameter: 1.5cm) to a wide rectangular target (width: 10cm; depth: 1cm), placed 15 cm apart (FIG 1). Since the main goal of this experiment was to assess the influence of reward on motor decisions during ongoing movements, we designed an experimental setup in which the influence of motor cost, on the choices, was only marginal. To this end, we rotated the geometrical arrangement 135 degrees’ counter-clockwise, so as to align a potential movement towards the centre of the rectangle with the direction of maximum arm inertia (FIG 1C). In other words, movements towards either side of the rectangle implied the same motor cost (Cos et al. 2011). The task contained two kinds of trials: baseline and change of mind (CoM) trials. The subject was instructed to make a reaching movement from the origin cue to a freely selected position along the length of the rectangular target, while attempting to maximize reward. The reward obtained in each trial was contingent upon arrival position along the length of the rectangle and the final distribution of value (DoV). The DoVs were bimodal, peaking at the right and left along the length of the rectangle, and decreasing towards zero when approaching its centre. The distribution was also equal to zero off the right/left sides of the rectangle, meaning that a reaching movement missing the target, or entering from the sides would not yield any reward. The quantitative value of the reward was represented by the height of the peaks, which matched the reward points in cm. The participant was presented with one of three possible DoVs: 3-3, 1-5 or 5-1 (FIG. 1C).

**FIGURE 1.**
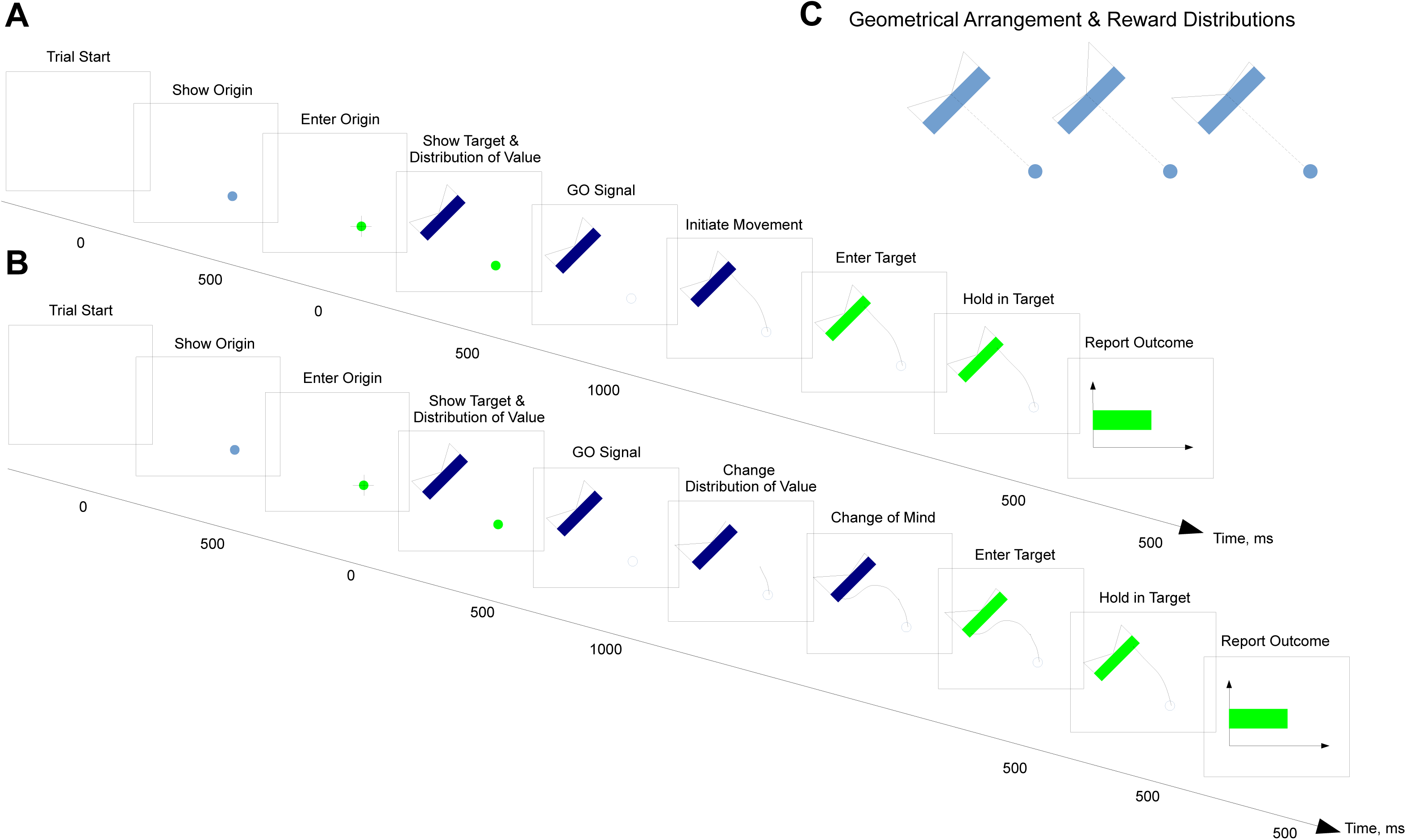
**A-B.** Both kinds of trials start with the presentation of a blank screen during 500ms. After this time, a 1cm diameter pale blue origin cue is presented on the bottom right part of the screen. 1ms after the subject’s fingertip crosses its border, the origin cue changes to green, and the recetangular target is presented on the top left part of the screen, 15cm away from the centre of the origin cue and rotated 135 degrees. The rectangle is 10cm wide and 1cm deep. Simultaneously, the distribution of value is presented as two rectangular triangles, centred at the midst of the rectangle wide side, and peaking on either side. One of three possible distributions was presenteed: 1-5, 3-3 or 5-1.The GO signal was given 100ms after the presentation of the distribution, by turning the origin cue to white (the background colour). For the CoM, some time after the movement onset, a second distribution of value was presented during 250ms. Target arrival was signaledd by turning the rectangle from blue to green when the end-point entered the target. A red horizontal bar provided visual feedback as to the precision of the movement. **C**. Geometrical arrangements associated to the reward distributions shown in this experiment: the distribution of value was always shown on top of the rectangle target. One of three distributions were presented: 3-3, 1-5, 5-1, shown from left to right, respectively.

Real-time, visual feedback of hand position was provided during the trial by a 1cm cross on the screen, synchronized with the tip of the participant’s right-hand position on the experimental table. The time-course of each trial is shown in FIG 1A-B. A trial began when the origin was shown on the screen and the subject entered the cue. Approximately 1000ms later, the rectangular target and the initial distribution of reward were shown (FIG 1A-B). After a 1000ms observation interval, a GO signal was given by turning the origin cue white. The subject was then instructed to perform his/her selected path trajectory towards the position along the rectangle, which they deemed most rewarding. If the subject left the origin before the GO signal was given, the experimental arrangement disappeared and the subject had to wait until the regular trial duration of 7s elapsed, before attempting again. Correct target entry resulted in the rectangle turning green. After 500ms of holding the position within the target, the reward associated with the specific motor path was shown for a duration of 500ms. The screen turned white after an interval that equalized the entire trial duration to its fixed duration of 7s. The CoM trials followed the same time-course of the baseline trials, with the exception that the distribution of value changed 100-250ms after movement onset.

Each participant performed a single experimental session consisting of N=630 trials (∽1h15min), divided into 7 blocks of 90 trials each. Each block consisted of 72 baseline trials (24 repetitions of each DoV) and 18 CoM trials. There were 18 types of CoMs trials, as a function of the change of DoV (3-3/1-5; 3-3/5-1; 1-5/3-3; 1-5/5-1; 5-1/3-3; 5-1/1-5) and the time at which the DoV change occurs (Early (E; t < 80ms), Medium (M; 80ms < t < 140ms) or Late (L; t > 140ms ± 30ms)) --- TABLE 1. Each block contained one trial of each possible CoM type. Trial order was counterbalanced and randomized both within and across blocks. The second DoV was shown for a duration of 300 ms.

Subjects were told that the DoV would change on some trials, and that, if they wanted, they could change their mind and adjust their initial motor path. Feedback was provided at the end of each trial in the form of a green horizontal bar, shown during 1000ms. The length of the bar was proportional to the reward obtained as a result of their reaching movement. The inter-trial interval was modulated to maintain a fixed 7s trial duration, to prevent participants from increasing their speed to maximize reward.

### Analysis of Kinematics

Analysis of the processes underlying CoM was carried out through careful examination of motor trajectories and their associated velocities and accelerations. We used two main metrics to characterize the processes related to changes of mind: (1) the time from the presentation of the second DoV to the hard bend of the trajectory indicating the change of mind (tCoM; FIG 3A), and (2) the probability of changing your mind (PCoM). Trials were discarded if the response time was longer than the timeout of 7s, if the subject entered the target rectangle through the sides or top and/or if he/she left the origin before the GO signal. To characterize the dynamics of CoM, we also analysed the sequence of kinematic metrics typical of reaching movements (FIG 3C). On a single trial basis, we calculated the reaction time (RT) and identified the following kinematic markers from the tangential velocity: peak acceleration (PA), time-to-peak acceleration (TTPA), peak velocity (PV), time-to-peak velocity (TTPV), peak deceleration (PD), time-to-decelerate (TDEC), time-to-peak-deceleration (TTPD) and overall movement time (MT). On CoM trials, we also considered kinematic markers of the radial velocity after the CoM, namely the peak radial velocity (PRV), the time-to-peak radial velocity (TTPRV), the overall switch movement time from the trajectory hard bend to the movement offset (MTCoM), the time of deceleration, from the TTPRV to the movement offset (TTPDECCoM) --- FIG 3C.

**FIGURE 2.**
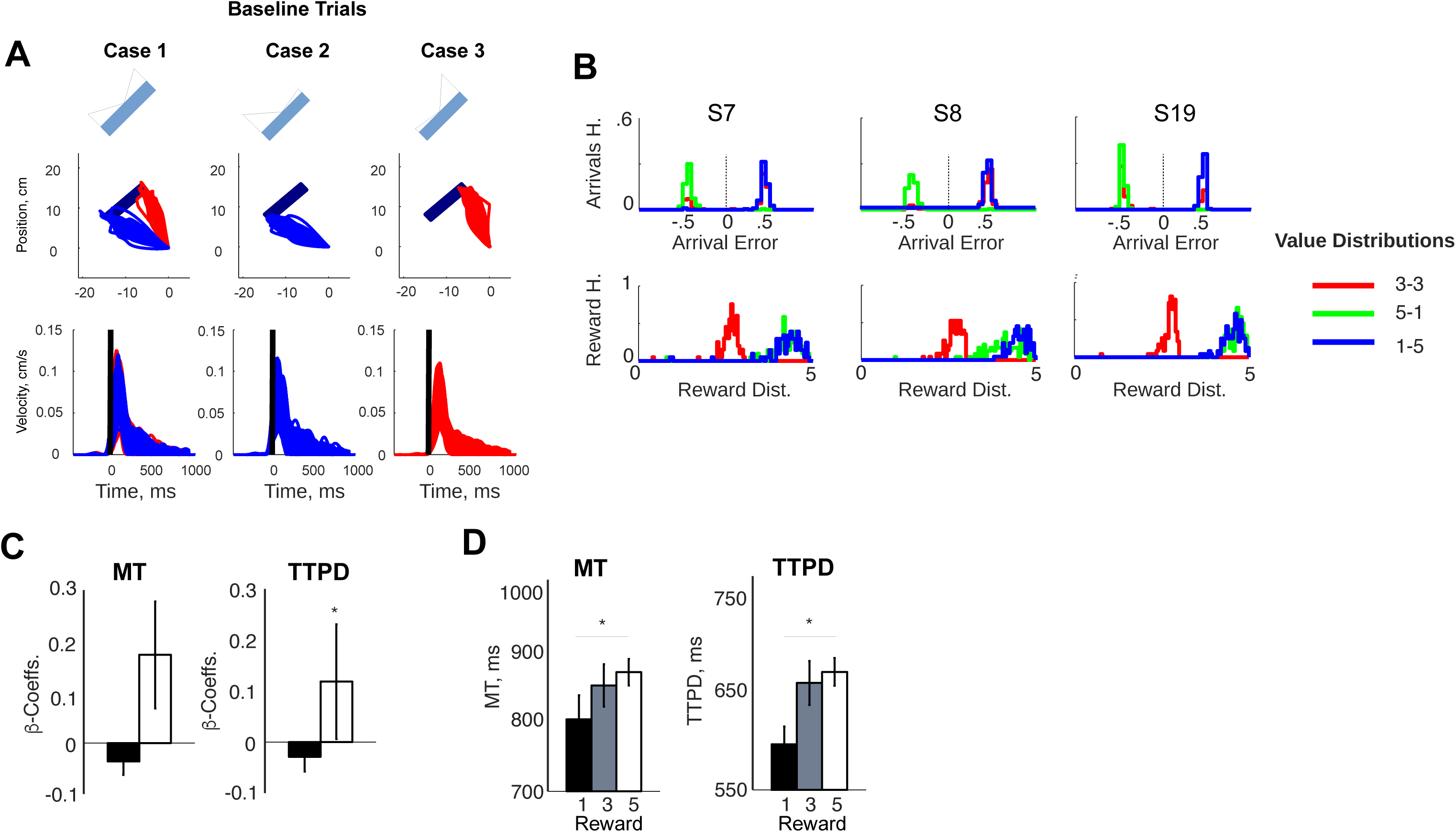
**A.** Top: Baseline trajectories for a typical subject, plotted for each distribution of reward value: 33, 51 and 15, colour coded as a function of their selected side of the target (right: red; blue: left). Bottom: baseline tangential velocities for each distribution of reward value: 3-3, 5-1 and 1-5. **B.** Arrival error/reward (top/bottom) distributions for three typical subjects (S7, S8, S19), colour coded as described in A. **C**. Grand average b-coefficients for the GLMs for the Movement Time (MT) and Time-to-Peak-Deceleration (TTPD) as a function of Reward Value (V). **D.** Mean and average MT and TTPD as a function of Reward Value.

**FIGURE 3.**
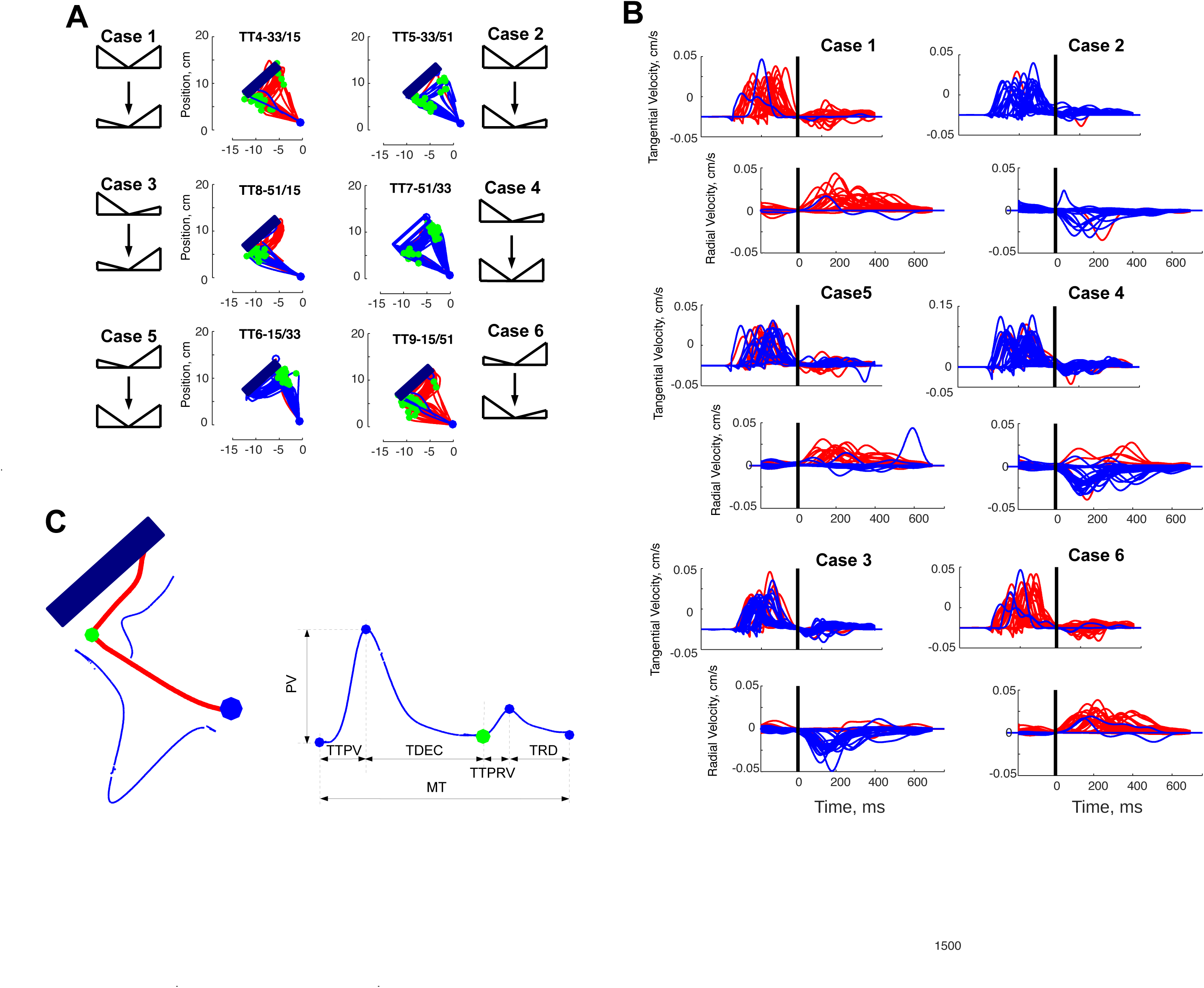
**A.** Change of mind trajectories for the six types of CoM trial considered (1. 33/15; 2. 33/51; 3. 15/33; 4. 51/33; 5. 51/15; 6. 15/51), for trials in which the right/left (red/blue) side of the rectangle has been aimed for. **B.** Change of mind tangential and radial velocities for S9, at each of the six aforementioned cases, aligned at the moment in which the trajectory bends hard to the side opposite to their initial choice. **C.** Kinematic Markers for the tangential (PV, TTPV, TDEC) and radial (TTPRD, TRD) velocities of a CoM trial, and overall Movement Time (MT).

### Generalized Linear Models for Statistical Analysis

Statistical analyses were carried out to assess how the value of a prospective reward influences the metrics of movement and the probability of changing your mind (PCoM). To this end, we fitted a series of Generalized Linear Models (GLM) to each of the aforementioned kinematic and behavioral metrics, for each individual participant. For the analysis of baseline trials, the GLMs were regressed against a single factor: the value of the reward intended by each trajectory (V) --- FIG 2C.

The goal of the second set of analyses was to determine the influence of factors related to CoM on movement parameters and on the likelihood of switching target side. Thus, we operationally defined the notion of prospect gain (G) as the difference in value between the current and the alternative choice, after the second DoV was presented. In other words, G quantifies the desirability of the CoM as compared to the possibility of continuing the original reaching movement. For the analysis of the tCoM (FIG 5), we performed a second set of GLMs whose independent factors were: the Gain (G) to be attained if the change of mind were carried out, the time of presentation of the second DoV from movement onset (T), and the tangential Peak Velocity (PV). Statistical group significance was established via *F*- and *t*-tests on the regression coefficients for each factor obtained per participant.

**FIGURE 4.**
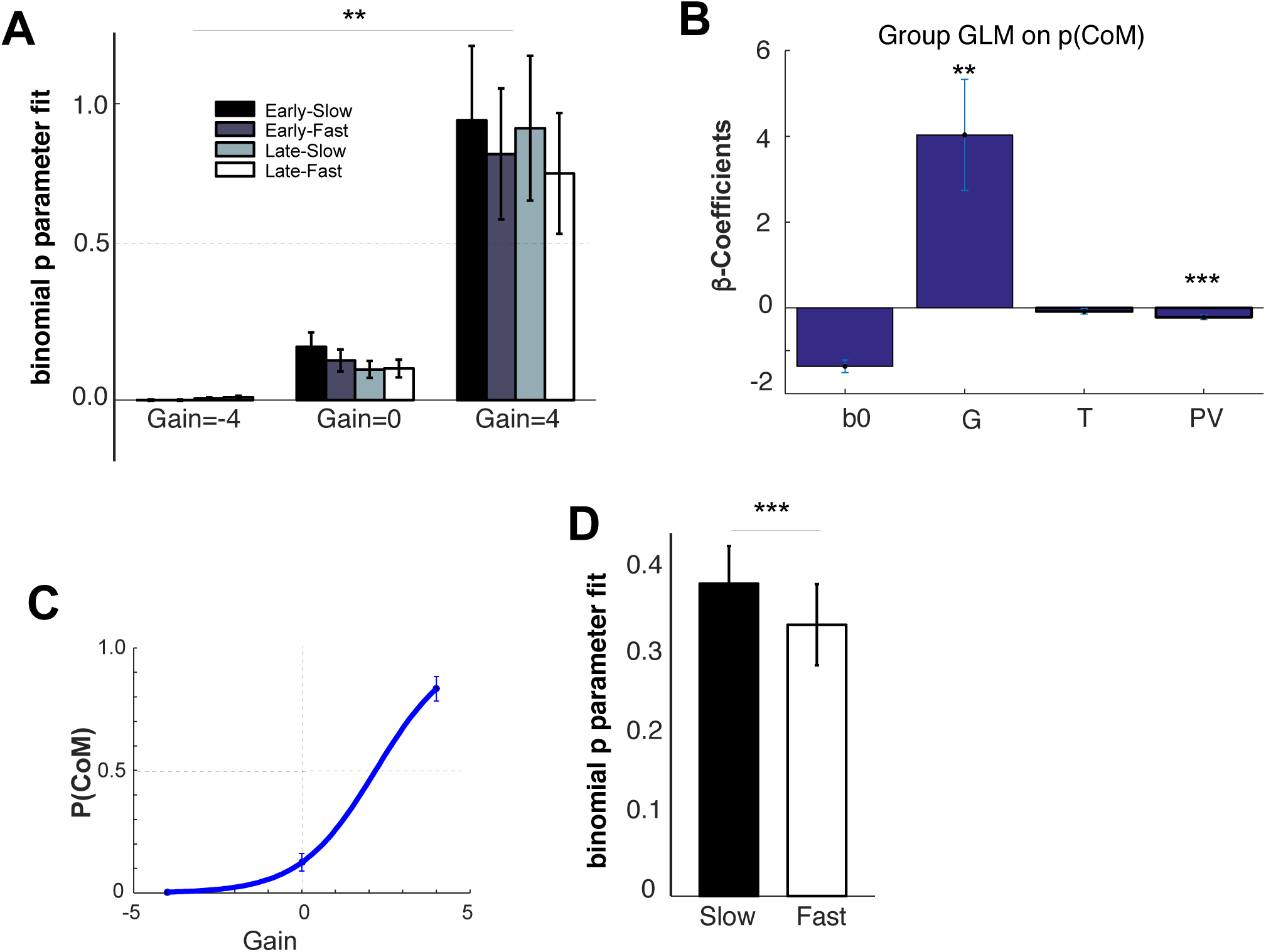
**A**. Group average binomial parameter p fit for the PCoM, as a function of Reward Gain, Time of presentation of the second distribution of value, and peak tangential velocity for the associated movement. Mean and standard errors for each case are plotted. **B.** Grand Average Regression coefficients of GLMs performed on the binomial p parameter fitted to the Probability of changing target (PCoM), as a function of three factors: reward gain if CoM (G), time of presentation of the second distribution of value (T), and the peak tangential velocity (PV). The PCoM exhibits a main increasing effect with the prospect of Gain (F(15,3)=9.71, p=0.0082) and a decreasing effect with velocity (F(15,3)=15.79, p=0.0016) The time of presentation of the second distribution exerts no significant effect on the PCoM F(15,2)=1.88, p=0.21. **C.** Group Average effect of Reward Gain onto PCoM (binomial fit p-parameter). **D.** Group Average velocity Effect on PCoM (binomial fit p-parameter).

**FIGURE 5.**
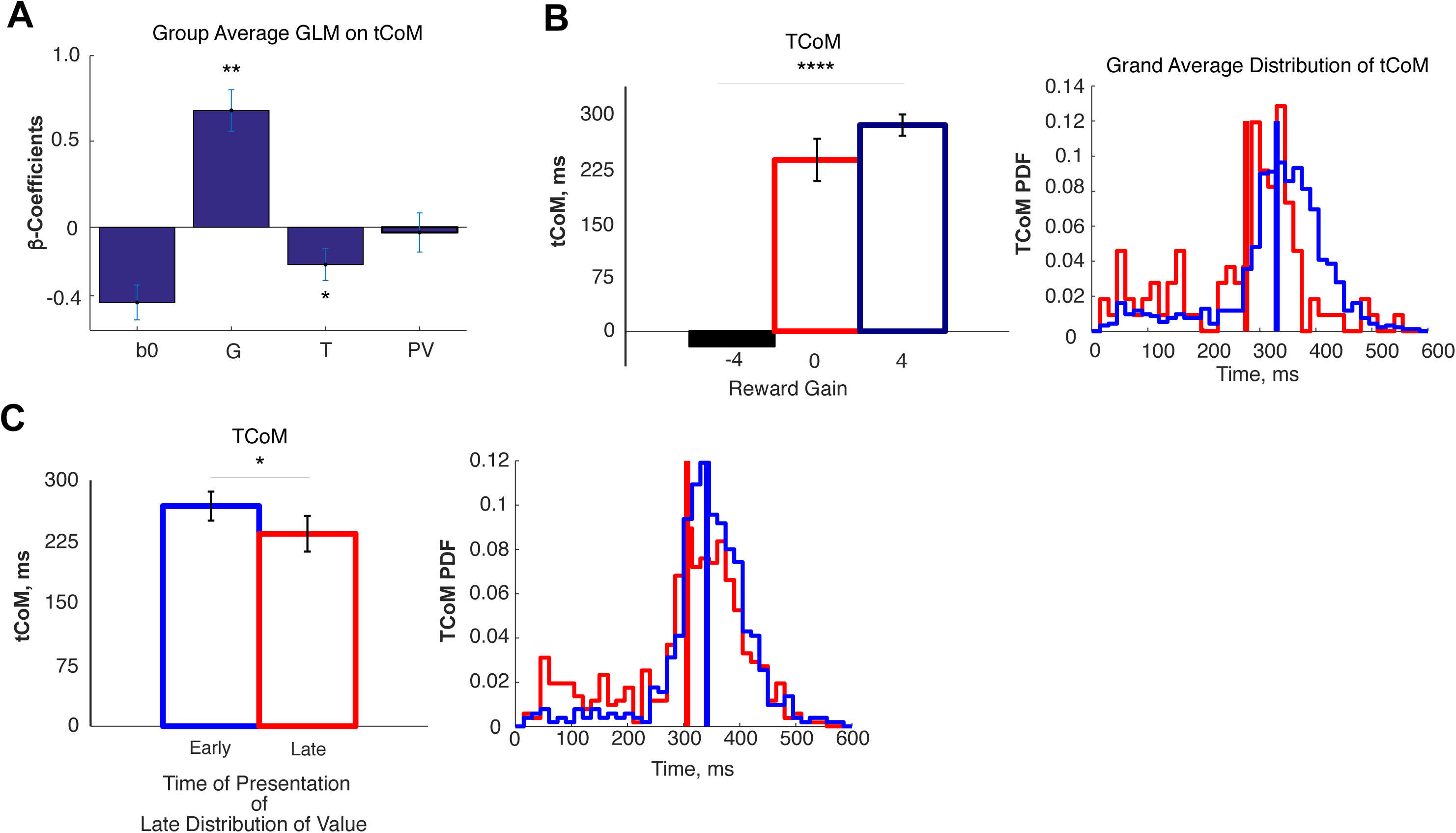
**A.** Grand average regression coefficients of GLMs performed on the time of Change of Mind (tCoM), as a function of three factors: reward gain if CoM (G), time of presentation of the second distribution of value (T), and the peak tangential velocity (PV). The tCoM exhibits a main increasing effect with the prospect of Gain (F(15,3)=31.82, p=0.0001) and a decreasing effect with the time of presentation of the second distribution of value (F(15,3)=5.4571, p=0.035. The movement velocity exerts no significant effect on the tCoM F(15,3)=0.0716, p=0.79). **B**. Left: Group Average effect of Reward Gain onto tCoM. Right: Group average tCoM histogram as a function of Reward Gain (Red: G<4; mean=247.956ms, **?**=51.62.ms/ Blue: G=4; mean=317.00ms, **?**=116.55.ms). A post-hoc t-test yields a statistically significant difference between distributions at p=2.98e-09. **C.** Left: Group Average velocity Effecf on tCoM. Right: Group average tCoM histogram as a function of the time of presenttion of the distribution (Read: Late; mean=27302ms, **?**=145.42.ms; Blue: Early; mean=305.75ms, **?**=133.40ms). A post-hoc t-test yields a statistically significant difference between distributions at p=7.46e-09.

The third set of analyses aimed at characterizing the unfolding dynamics of CoM trials. To that end, we divided the trajectories into two intervals: from movement onset until the hard bend, and from the hard bend until movement offset (FIG 3A). Note that this distinction was operationally possible because, at the switch, the participants’ tangential and radial velocities were zero on the trials where a CoM occurred (FIG 3B). We performed two sets of GLMs, one for the kinematic metrics calculated at each of the two segments. First, to assess the influence of the trial’s initial DoV, the first set used the following factors: the reward value of the initial choice (V), the time of presentation of the second DoV (T), and a binary variable indicating whether the participant changed their mind on that trial (CoM variable) --- FIG 6. Second, to assess the influence of the second DoV on movement dynamics, we regressed the same kinematic metrics against the prospect reward Gain (G), the time of presentation of the second DoV (T), and the binary variable indicating whether the participant changed their mind on that trial (CoM) --- FIG 7. To establish significance at a group level, *F*- and *t*-tests were performed on the regression coefficients associated to each factor across subjects.

**FIGURE 6.**
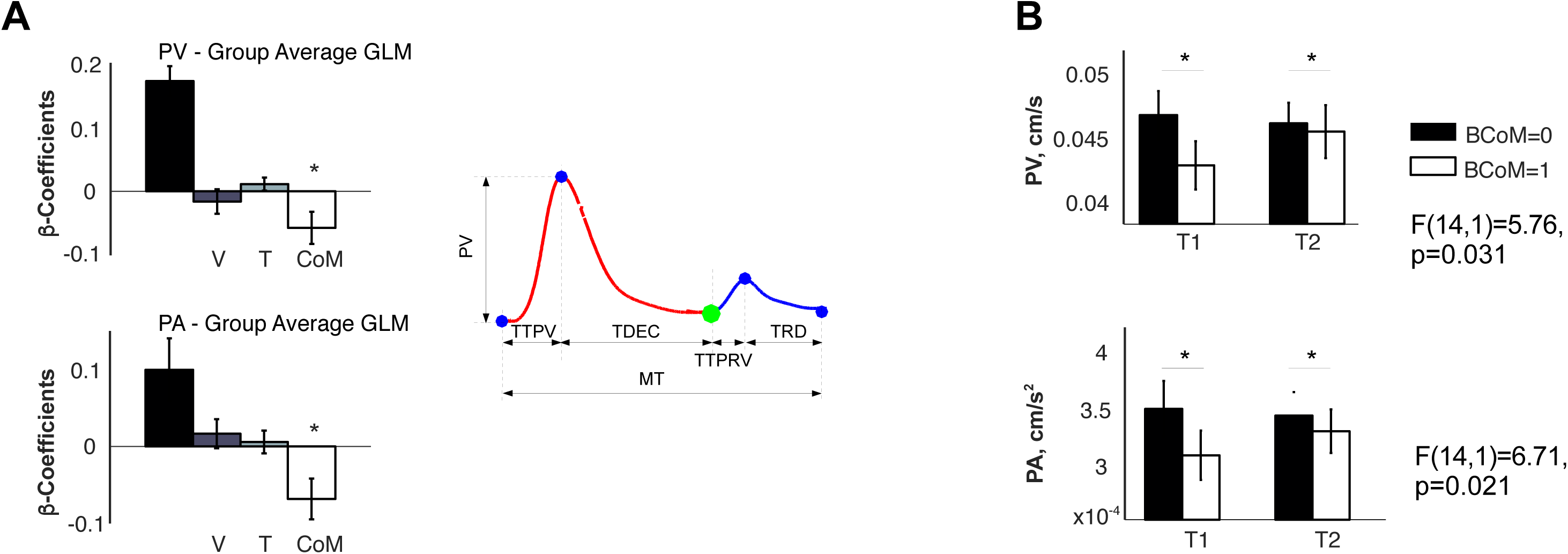
**A.** Grand average regression coefficients of GLMs performed on the Peak Velocity (PV) and on the Peak Acceleration (PA), as a function of three factors: reward value (V), time of presentation of the second distribution of value (T), and a binary variable indicating whether there was a CoM in that trial. Remarkably, both the PA and PV were sensitive to the CoM F(15,1)=, p=), but not to reward value or to the time of presentation of the second DoV. **B**. PV and PA as a function whether there was a CoM or not, for each Target.

**FIGURE 7.**
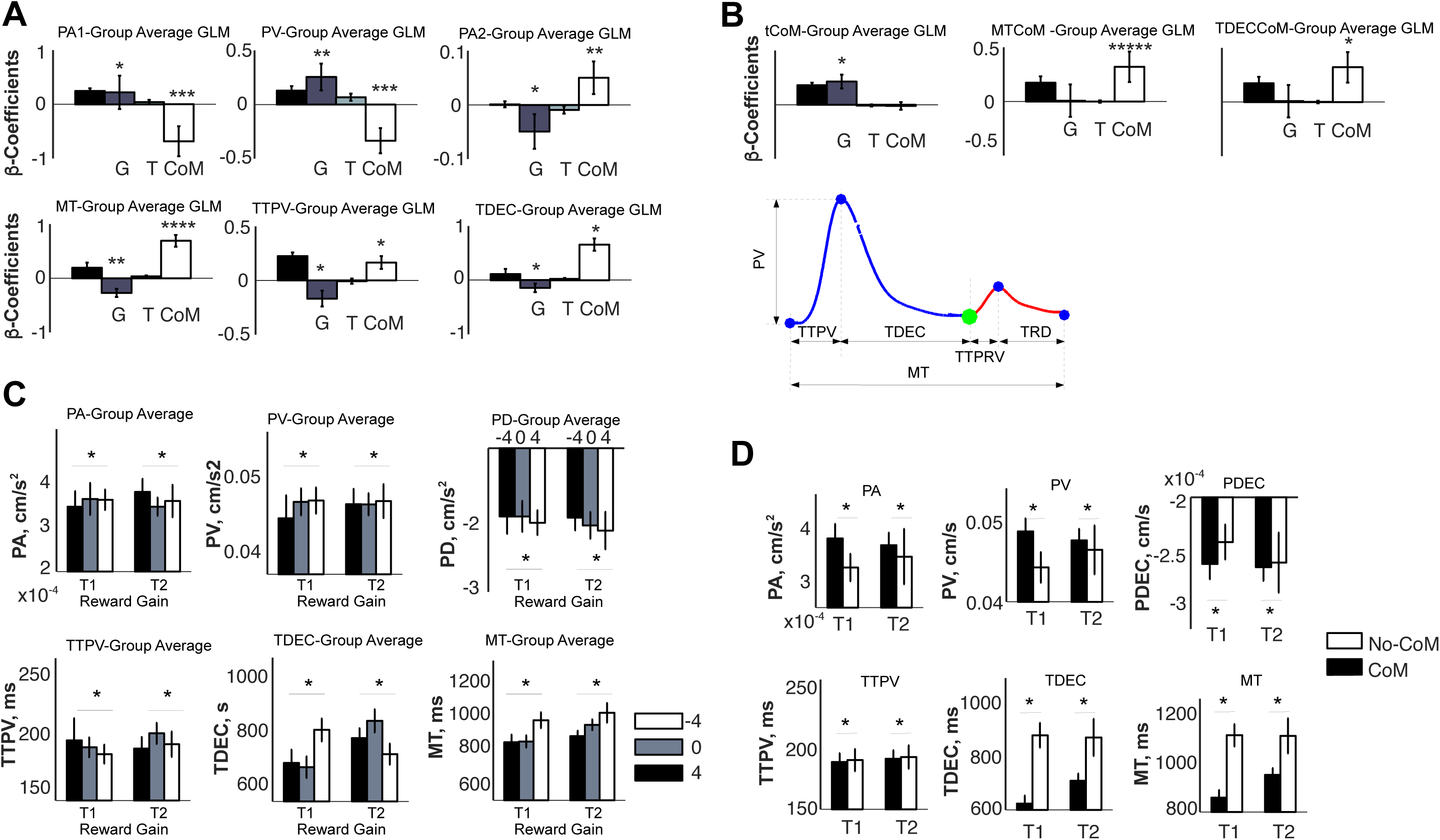
**A.** Grand average regression coefficients of GLMs performed on the Peak Acceleration (PA) and on the Peak Velocity (PV), Peak Deceleration (PD), Movement Time (MT), Time-To-Peak Velocity (TTPV), Time to Peak Deceleration (TTPD) as a function of three factors: reward gain (G), time of presentation of the second distribution of value (T), and a binary variable indicating whether there was a CoM in that trial. Remarkably, all six variables signal a significant difference between trials in which there was a CoM vs. those in which subject stuck to their original choice. **B**. Grand average regression coefficients of GLMs performed on the time of CoM (tCoM), on the Movement Time after tCoM (MTCoM), and on the deceleration interval (TTPDCoM) of the second segment of the trajectory. **C.** Group means and standard errors across subjects for the PA, PV, PD, TTPV, TDEC, MT, as a function of Gain. **D.** Group means and standard errors across subjects for the PA, PV, PD, TTPV, TDEC, MT, as a function of whether there was a CoM or not.

### The Probability of Change of Mind (PCoM)

To assess the probability of changing your mind (PCoM; FIG 4), we first calculated the amount of times subjects changed their mind by analyzing the subjects’ trajectories and then fit a binomial distribution for the proportion of times CoM occurred using the binofit function provided by MATLAB (The Mathworks, Natick, MA). The binomial distribution is characterized by a single *p*-parameter, which captures the likelihood of a binary event to occur. The fit was performed for each individual participant, and for each possible value of the prospect gain (G), for short and long presentation times of the second DoV (T), and according to the peak velocities (PV) of the behavioral response. Both T and PV were classified as Early/Late and Slow/Fast using median splits within the T and PV distributions of each individual participant. We then calculated a GLM of the resulting *p*-parameter of the binomial distribution against the same factors: G, T and PV for each individual subject. We established group significance by running an *F*-test on each of the regression coefficients. Post-hoc *t*-tests were also performed to confirm their significance. Finally, to plot the dependence of the PCoM on prospect Gain, we fitted a parametric sigmoidal curve to the *p*-parameter obtained as a function of each possible value of the prospect gain (G) at each individual subject, as shown by EQ 1.

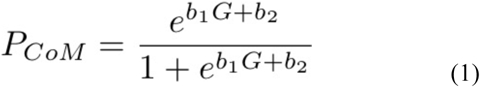

The quality of the fit was assessed with a determination index *r* and a *p*-value.

## RESULTS

### Baseline Preferences

FIG 2A shows a set of typical trajectories from the origin cue to a position along the length of the rectangular target, for all three baseline value distributions described in FIG 1C (3-3; 5-1; 1-5). Consistent with instruction, the trajectories show that this participant’s choices strongly favour the side of the rectangle offering the largest reward prospect for the 5-1 and 1-5 distributions and are more evenly distributed for the 3-3 distribution. These observations are also reinforced by the distribution of arrival positions and related rewards shown in FIG 2B for three typical participants (7, 8 and 19).

Tangential velocity profiles are shown in FIG 2A for all three baseline cases, aligned with movement onset. The profiles exhibit a fast rise to peak and a longer deceleration phase until target arrival, consistent with the need of a slow, controlled movement in the interval immediately preceding target arrival and subsequent reward delivery. This is confirmed by an F-Test across the regression coefficients of MT and TTPD against reward value for each participant (FIG 2C-D). The F-test shows that the duration of the TTPD interval (from peak velocity to movement offset) increases with value (*F*(1,15) = 6.21; *p* = 0.0248), and, although marginally non-significant, also the entire MT (*F*(1,15) = 4.5; *p* = 0.051). By contrast, the TTPV does not significantly vary with reward value (*F*(1,15) = 3.78; *p* = 0.0708). This effect is similar to Fitt’s law (Fitts, 1954) in sofar the subject is more concerned by the need for proper aiming than aroused by the prospect of gaining more reward (Dayan and Balleine 2002; Niv 2007; Prevost et al. 2010).

### Change of Mind Trajectories

Figure 3A shows a set of trajectories during Change of Mind (CoM) trials, parametrized as a function of their initial/final distributions of reward value (3-3/1-5; 3-3/5-1; 5-1/1-5; 5-1/3-3; 1-5/3-3; 1-5/5-1). The trajectories confirm that the subject’s ultimate goal is to gain the largest possible reward, given that target arrivals are most frequent at positions nearby the largest reward offer. However, we distinguish two major behavioral strategies to attain the desired arrival position. First, similar to behavior on baseline trials, one strategy would consist of initially reaching for the position offering the largest reward, and to alter that ongoing movement if the alternative side offers a better reward after the showing the second DoV. The second strategy, in DoVs where there is a strong imbalance (e.g., 1-5/5-1), consists of initiating a trajectory towards the least appealing side and changing the motor path if the second distribution confirms the presumed bad prospect of the initial movement. In a way, the first strategy assumes that the initial distribution will not change (there will be no second distribution), while the second one hopes for a change in the prospective value of the current distribution, as the movement progresses.

Figure 3B shows a set of typical tangential and radial velocities for all six aforementioned CoM trial cases. The velocities are aligned at the time of CoM (plotted in green, FIG 3A), which effectively partitions the movement into distinct phases. First, a ballistic movement, tangential from the origin cue, and second a radial movement towards the alternative target side.

### Changes of Mind during Ongoing Movements

We anticipated that participants would change their mind whenever the alternative option offered some gain with respect to their original choice. Consistent with this, our first observation of the trajectories in FIG 3A indicates that participants changed their trajectory most frequently when the second DoV revealed a better alternative. Furthermore, to test the potential influence of time and velocity on the pattern of CoM, we classified trials as Early/Late and Slow/Fast by performing mean splits on the distribution of the presentation times of the second DoV and on the distribution of Peak Velocities preceding the CoM, respectively. Next, we fitted a binomial distribution to the proportion of CoMs experimentally observed for each of the 6 Gain × 2 Times × 2 Velocities cases at a single participant level, obtaining separate *p*-parameter values for combinations of Gains, Times and Velocities (METHODS). The grand average of the binomial p-parameter as a function of Gain, Time and Velocity, across subjects, is shown in FIG 4A.

To establish the statistical dependence of the Probability of Change of Mind (PCoM), we performed a GLM on the set of binomial p-parameter values for each case and participant, as a function of three experimental factors: the reward Gain (G) associated with the CoM, the time (Early/Late) of presentation of the second DoV, and the peak velocity (Slow/Fast) prior to the tCoM. The grand average regression coefficients obtained is shown in FIG 4B. Consistent with our hypothesis, an F-test of the beta coefficients for G across subject confirmed a significant and increasing effect of Gain on PCoM (*F(*15,3) = 9.71, *p* = 0.0082), reinforced by a regression of PCoM as a function of Gain, as plotted in FIG 4C. Remarkably, the same F-test on the coefficients of Velocity yields a significant decreasing effect on PCoM (*F(*15,3) = 15.79, *p* = 0.0016), meaning that the faster is the movement, the less likely is the CoM (FIG 4D). An F-test on the time of presentation of the second distribution of value (T), yields a non-significant effect on PCoM (*F(*15,1) = 1.88, *p* = 0.21).

### The Time to Change your Mind

In addition to the probability of altering the initial motor trajectory, we also calculated the time of CoM (tCoM) for every CoM trial and trajectory. The tCoM is defined as the time interval between the presentation onset of the second DoV and the hard bend of the trajectory, indicating a commitment to the change of target side. Consistent with the participant’s intent to better control arrival into the target, thus maximizing reward, subjects displayed increased MTs and TTPDs with increasing value on baseline trials. It would be fitting to hypothesize that the value to be gained by changing target side as well as the time of presentation of the second DoV are influential on the subjects’ urge to adjust their motor trajectory, and consequently tCoM. To test this, we performed a GLM of the tCoM for each individual subject as a function of three factors: the gain (G) associated with the executed CoM, the time of presentation of the second DoV, and the peak tangential velocity. FIG 5A shows the grand average across subjects of the regression coefficients for that GLM. A subsequent F-test on each factor coefficient yielded two group effects: a positive effect of Gain on tCoM (F(15,1) = 31.82, *p* = 0.0001), and a decreasing effect as a function of the presentation time of the second DoV (tDoV) (*F(*15,1) = 5.4571, *p* = 0.035). In other words, the larger the Gain, the slower the CoM; and the later the tDoV, the faster the CoM. Interestingly, although velocity is influential on the PCoM, it does not exert a significant effect on the tCoM (*F(*15,1) = 0.0716, p = 0.79). FIG 5B shows the grand average of the tCoM across participants as a function of Gain, as well as the histograms of the tCoM for the cases of G <= 0 and G = 4. FIG 5C shows the grand average of the tCoM as a function of the tDoV (Early/Late) and the distribution of tCoM as a function of the tDoV.

### Early vs. Late Change of Mind Kinematics

To analyze the dynamics of the adjustments in motor trajectories represenative of CoM, we partitioned CoM trajectories into pre-CoM and post-CoM intervals. The pre- and post-CoM segments were segmented according to the instant post movement onset at which both the tangential and radial velocity were equal to zero (FIG 3B). A zero-value indicated a hard bend in the initial trajectory towards the opposite side of the target from their original choice. Although we assumed that the trajectory bend signalled the switch, we hypothesised that the preparation for the CoM began well in advance. Thus, to test this prediction we observed the pre-CoM interval and assessed the influence of the reward value (V) the participant initially aimed for, based on the first DoV, the time of presentation of that DoV, and a binary variable indicating whether a CoM occurred. We regressed the RT, PA, PV, TTPA, TTPV, PD, TTPD on each of these three variables.. FIG 6A-B shows that the only two metrics exhibited a significant decreasing effect as a function of CoM: the peak acceleration and peak velocity (PA; F(14,1) = 6.71, p = 0.021; PV; F(14,1) = 5.76, p = 0.031). Indeed, although the driving force to change the initial trajectory may be the reward on the converse side of the target, these results show that the initial state of the motor system, characterized by the initial peak acceleration, is one that is prone to change if the reward gain is beneficial.

In a complementary fashion, we performed a set of analyses to determine the influence of the second DoV on movement kinematics for the pre-and post-CoM kinematic markers. Specifically, rather than the reward associated with their initial choice, our regression factors on movement kinematics were: the reward Gain (G) that would result from switching to the other side of the target versus that of staying with the current movement, the presentation time of the second DoV (T), and the binary variable CoM, which indicated whether or not participants changed their mind on each trial. We regressed GLMs for each participant as a function of these three factors (G, tDoV, CoM) on to two sets of kinematic variables: those preceding the CoM (PA, PD, PV, TTPV, TDEC, tCoM, MT), and those following it (MTCoM, TTPDCoM) --- see Methods & FIG 7D. FIG 7A shows the grand average regression coefficients for the variables preceding the trajectory bent, while FIG 7B shows those succeeding the CoM. It is important to note that the F-tests across subject coefficients for each GLM factor report two significant main group effects. First, significant differences were found between CoM and non-CoM trials (CoM factor). In particular, these differences were seen in increased durations of MT (F(15,1) = 31.76, p = 0.0001), TTPV (F(15,1) = 9.27, p = 0.0094), MTCoM (F(15,1) = 39.17, *p* < 1E-5) and TTDEC (F(15,1) = 24.82, *p* = 0.0003); as well as increased amplitudes of PA2 (*F*(15,1) = 9.68, *p* = 0.0083). Furthermore, decreased amplitudes of PV (*F*(15,1) = 15.73, *p* = 0.0016) and PA (*F*(15,1) = 10.75, *p* = 0.0060); as well as a decrease in duration of TTPDCoM (*F*(15,1) = 6.74, *p* = 0.022) were reported. Second, a Gain effect, elongating TTPV (*F*(15,1) = 9.36, *p* = 0.0091), TTPD (*F*(15,1) = 4.94, *p* = 0.044) and MT (*F*(15,1) = 10.97, *p* = 0.0056), as well as increasing the amplitude of PD (*F*(15,1) = 8.42, *p* = 0.012), and decreasing the amplitudes of PD (*F*(15,1) = 10.52, *p* = 0.026) and PV (*F*(15,1) = 10.52, *p* = 0.0064) respectively, were found FIG 7C-D. In conclusion, these results would strongly suggest that although the bend in trajectory occurs roughly 300ms after the second DoV appearance (FIG 5), the decision to switch motor plans occurs and is prepared much sooner, from about 100 ms after the presentation of the second DoV (TTPA).

## DISCUSSION

The goal of this study was to investigate how changes in reward prospect influence decision-making once commitment for an option has been made. Although previous evidence has shown that activity in the pre-motor cortex may encode several actions simultaneously (Baumann et al. 2009; Cisek and Kalaska 2010; McPeek and Keller 2002), reasonable doubt remains as to whether the simultaneous encoding of several motor actions extends to the execution phase, well after a specific option has been selected and the movement is ongoing. Here, we assumed that decisions abide by a principle of cost-benefit optimization where the option with the highest prospect is most often selected, assuming that all other factors remain equivalent. Furthermore, we hypothesized that a change of reward prospect at any point in time should dynamically drive adjustments in the desirability of each option. This, in turn, would necessitate a rapid reassessment of costs/benefits across prefrontal and motor cortical areas, ultimately resulting in a change of mind and ensuing motor trajectory. To test this hypothesis, we performed a set of experiments in which human participants were instructed to freely select a reaching path-trajectory from an origin to a wide rectangular target. The quantity of the reward was contingent upon the distribution of reward along the rectangle side and precision at the arrival point. To test our hypotheses, reward distributions were changed in one third of the trials post movement onset. Our results showed that participants were most likely to alter their initially selected motor plan when the value of the end reward along a different path was better, even during an ongoing movement. Furthermore, the changes of mind occurred more frequently during slow movements, and required a duration on average inferior to the reaction time. Finally, the short latency of the CoMs, along with the fact that the first time-to-peak-acceleration exhibit significant differences during CoM and non-CoM trials, strongly suggests that the decision to adjust one’s trajectory occurs as early as the first peak acceleration.

### Baseline Effect of Reward

First, we examined the influence of rewad value on kinematic parameters during baseline trials. Several studies have shown that larger rewards tend to increase movement vigour (Miyachi et al. 2002; Watanabe 2007), its energy (Weiner & Joel, 2002) and frequency (Jackson, Anden & Dahlstrom, 1975). In contrast, our results are consistent with a concern for potential loss, expressed through an increase of the overall movement time and of the duration of the deceleration phase (FIG 2D). Although this does not rule out a potential vigour effect, it does indicate that loss aversion is stronger when reward is at play and the prospect of losing it is significant. It is important to keep in mind that reward in this task is contingent upon precision: responses hitting the extremes of the rectangle length were awarded a value close to the maximum, while those that missed the rectangle, which may be very close to the peak, received zero. Thus, it would be adaptive to develop a behavior consistent with risk aversion when there is a large reward at stake (Wu et al. 2011).

### Gain and Noise to Change your Mind

This study aimed at evaluating how changes in reward distribution affect decision-making during the execution phase. Our results show that PCoM increased when the second DoV presents an opportunity for greater reward on the side opposite to our initial trajectory, and that the PCoM decreased when this opportunity is presented later in time. Remarkably, the PCoM amounts 10-15% in the absence of Gain (FIG 4A), implying that although most decisions aim at the largest reward prospect, some other times subjects also opt for a choice of lesser gain. Although this does not invalidate the main principle of seeking reward, this may be interpreted as an effect of neural noise, interfering with the commitment to a specific action in the presence of multiple options (Cisek and Kalaska, 2010).

In a similar fashion, tCoM exhibits the same increasing sensitivity to Gain as the PCoM, but it also decreases as a function of the presentation time of the second DoV and is insensitive to velocity. Consistent with the same hypothesis of neural noise, the mean tCoM is shorter when the Gain is zero than when the Gain is large (FIG 5). This would imply that CoMs whenever there is no reward to gain are either made in the absence of proper processing of value, or guided by the concern of losing the reward offered by the alternative side, and biased by neural noise. This is also consistent with the fact that CoMs often occur close by the presentation of the second DoV, rendering a hypothetical pre-frontal analysis of the second distribution of value unlikely. Moreover, this would be consistent with the fact that changing your mind when there is nothing to gain is counter-productive. Finally, tCoM occurs sooner when the presentation of the second DoV occurs later (FIG 5C), suggesting an increased urgency for change.

### Reward Value onto Ongoing Behavior

Our analyses have also shown that the manner in which CoMs are implemented during reaching movements is consistent with the interplay between two sequential movements. The first movement occurs from the origin to the offset of tangential velocity, and the second from the onset of radial velocity to its final offset. Importantly, the influence of the first DoV on movement is constrained to a magnification of the late deceleration (TTPD) with reward, consistently with a risk aversion effect. By contrast, the second DoV exerts a broad influence on the manner in which participants move towards the target by extending overall movement duration and weakening movement intensity (most kinematic markers are influenced by reward Gain --- PA, PV, PD, MT, TTPV, TDEC, tCoM), both during the first and second movement segments (FIGs 6-7), shaping both the acceleration, deceleration phase and the specific manners in which the target is entered. Furthermore, our analyses of kinematics have also pointed out that as early as at the first PA, the movement is influenced by the CoM. This implies that, although the hard trajectory bend occurs on average 300ms after the presentation of the second DoV, the conditions necessary for the change of mind are already established around the first PA (80-90 ms), a few ms after the second DoV has been presented. The very short latencies obtained strongly support the notion that several motor plans are simultaneously encoded by pre-motor cortical areas (Cisek 2007; Cisek and Kalaska 2005, 2010; Cos et al. 2011), and that some decisions in these experiments were likely made without full consideration of the presented values. We believe that this very short latency may have been facilitated by the visual coding of reward we used in our experiment, which is known to facilitate the processing of visual stimuli, as size is the only element that matters to assign reward value in our task.

### Conclusion

The results from the analyses on PCoM and tCoM support the notion that reward exerts an influence not only on the choice itself but also on how movement are executed. Previous studies have pointed to the role of goal-directed behavior when planning and executing motor decisions (Cisek, 2007; Nashed et al., 2012), which is also supported by this study. Participants changed their minds and adapted their trajectories based on target value and these adjustments were modulated post-movement onset. Furthermore, value appeared to exert an influence on online feedback processes in several ways. First, there was a modulation of the time it takes to reprocess a reward (tCoM) as a function of the interplay between the presented value case and the time of DoV change. Thus, we change our mind if given sufficient time, and if the reward associated to an alternative second option outweighs the current one. Second, in the context of voluntary movements, the motor system not only takes into consideration a variety of environmental factors and intrinsic biomechanical and external costs (Cisek and Kalaska 2010; Cos et al. 2012; Nashed et al. 2014), but also the perceived (cognitive) value of the target, supporting the notion that decision-making models should factor in implementations of cost/benefit trade-offs like utility. Third, given the fact that changes of mind do occur on average quicker (∽300ms) than the RT prior to movement onset (∽400ms), this supports the parallel planning hypothesis, given that these adjustments must be made in a relatively small time-frame and require a rapid response. Fourth, feedback corrections do in fact appear to share the sophistication of the motor system for planning and executing motor actions. Thus, if the new distribution of value is desirable (larger value) and there is enough time to adjust, then we are likely to change our mind.

### Limitations and Future Directions

As a function of the results obtained in this study, there are some aspects that remain to be addressed. First, it would be interesting to look into the risk aspects of behavior (aversive or not), to determine whether and how these exert a significant influence on kinematic parameters and changes of mind in response to changing value. Finally, given that the results of this study do not control for saliency (are we reacting to bigger reward or simply ‘bigger’ target size), it would be important to run an inverse version of this task to determine that value is in fact the modulating variable.

## Acknowledgements

IC was funded by the Marie Sklodowska-Curie Research Grant Scheme, grant number is: IF-656262.

